# Benchmarking large language models for cell-free RNA diagnostic biomarker discovery

**DOI:** 10.1101/2025.08.20.671358

**Authors:** Hunter A. Gaudio, Andrew Bliss, Conor J. Loy, Daniel Eweis-LaBolle, Anne E. Gardella, Iwijn De Vlaminck

**Affiliations:** Meinig School of Biomedical Engineering, Cornell University, Ithaca, NY 14850

## Abstract

Large-language models (LLMs) can parse vast amounts of data and generate executable code, positioning them as promising tools for the development of biomarkers and classifiers from high-throughput omics data. Here, we benchmarked six LLMs, OpenAI’s o3 and GPT-4o, Anthropic’s Claude Opus 4 and Claude 3.7 Sonnet, and Google’s Gemini 2.5 Pro and Gemini 2.0 Flash, for disease classification based on plasma cell-free RNA (cfRNA) profiles obtained by RNA sequencing. We analyzed data from cohorts of children with Kawasaki disease (KD) or multisystem inflammatory syndrome in children (MIS-C), adults with active tuberculosis (TB) or other non-TB respiratory conditions, and individuals with myalgic encephalomyelitis/chronic fatigue syndrome (ME/CFS) or sedentary lifestyle. We assessed two tasks: (i) gene-panel design, where each LLM mined public knowledge to nominate diagnostic genes for use in machine learning (ML), and (ii) end-to-end modeling, where LLMs built an ML workflow directly from raw RNA-seq counts. In the first task, the LLM-derived panels captured canonical immune pathways and outperformed randomly selected genes in all cohorts. They underperformed panels chosen by differential gene expression (DGE) analysis in the KD vs. MIS-C and ME/CFS cohorts but performed comparably or better for the TB cohort. In the second task, o3 produced classifiers for KD vs. MIS-C that performed just as well as conventional statistical methods without human intervention. Performance for TB and ME/CFS cohorts was slightly lower than the conventional approach. These findings delineate current capabilities and limitations of LLMs in diagnostics and open a path for their future use in biomarker discovery.

## INTRODUCTION

Fewer than one percent of published biomarkers receive US Food and Drug Administration approval, in part because biomarker development remains complex and poorly standardized^1–5^. Therefore, robust, sensitive, and streamlined methods that can integrate existing biological knowledge are needed^6^. Plasma cell-free RNA (cfRNA) profiles offer a rich, minimally invasive readout of tissue injury and disease and show promise across diverse conditions for biomarker discovery^7–18^. Yet, converting cfRNA profiles into clinically useful signatures remains technically challenging, error prone, and time-consuming.

Large language models (LLMs) can capture vast biomedical knowledge and can generate executable code, positioning them to automate and potentially improve each step of this process^19,20^. Early successes have been reported in the use of diagnostic LLM’s for radiology and pathology^21,22^, and first attempts at LLM-guided omics analysis are emerging^23–26^. Nevertheless, systematic testing in omics-driven biomarker discovery remains limited.

Here, we test whether state-of-the-art LLMs can assist or even outperform purely statistical approaches in developing cfRNA-based diagnostic classifiers. We evaluate LLMs from OpenAI (o3 (newer) and GPT-4o (older)), Anthropic (Claude Opus 4 and Claude 3.7 Sonnet), and Google (Gemini 2.5 Pro and Gemini 2.0 Flash) across three clinical cohorts. First, we compare classifiers built from LLM-nominated gene panels with those built from randomly selected panels and from panels selected via differential gene expression (DGE) analysis. Second, we task each model with end-to-end construction of ML classifiers directly from cfRNA gene count data, encompassing feature selection, classifier construction, parameter tuning, and performance reporting. These experiments pinpoint where today’s LLMs add value and outline a roadmap for deploying LLMs in diagnostic biomarker development.

## RESULTS

We assessed six state-of-the-art LLMs (OpenAI’s o3 and GPT-4o, Anthropic’s Claude Opus 4 and Claude 3.7 Sonnet, and Google’s Gemini 2.5 Pro and Gemini 2.0 Flash) across three clinical cohorts that span a gradient of diagnostic difficulty: Kawasaki disease (KD) (n = 115) vs. multisystem inflammatory syndrome in children (MIS-C) (n = 50) (mean sequencing depth: 28.1M reads per sample)^7,27^, tuberculosis (TB) (n = 142) vs. symptomatic controls (n = 109) (31.7M reads per sample)^9,28–30^, and myalgic encephalomyelitis/chronic fatigue syndrome (ME/CFS) (n = 93) vs. sedentary controls (n = 75) (40.8M reads per sample)^11,31^ (**Fig. 1A**). KD and MIS-C represent closely related inflammatory conditions with overlapping clinical presentations but distinct pathophysiology: KD is an acute vasculitis of unknown etiology that primarily affects children younger than five years, whereas MIS-C is a post-SARS-CoV-2 hyper-inflammatory syndrome that affects older children and shows greater myocardial and gastrointestinal involvement^32–35^. TB presents another diagnostic challenge, with global detection gaps leaving millions of cases undiagnosed annually; the limitations of sputum-based diagnostics and the need to differentiate active TB from other lung diseases make this an important test case for novel biomarker approaches^36,37^. ME/CFS poses the greatest challenge, being defined solely by symptom criteria with no validated biomarkers; rigorous selection of sedentary controls is therefore essential to avoid spurious associations^38–40^.

**Figure 1:**
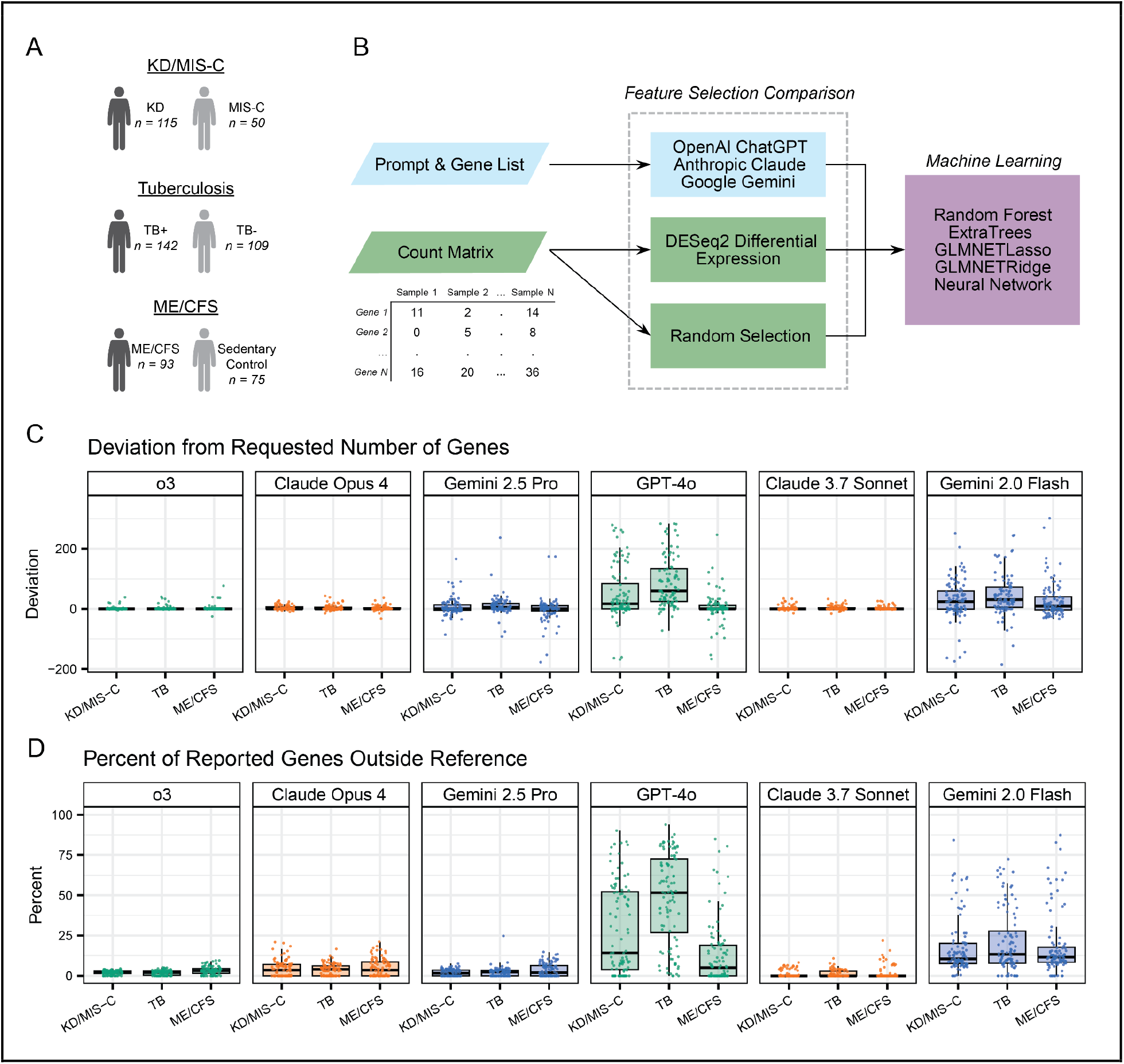
Sample/workflow overview and LLM adherence to prompt requests. (A) Patient cohort summary, (B) feature selection and ML comparison workflow, (C) deviation from number of requested genes (200) in LLM output (“long” input prompt), and (D) percent of LLM output genes not contained in the reference list (“long” input prompt).

**Figure 2:**
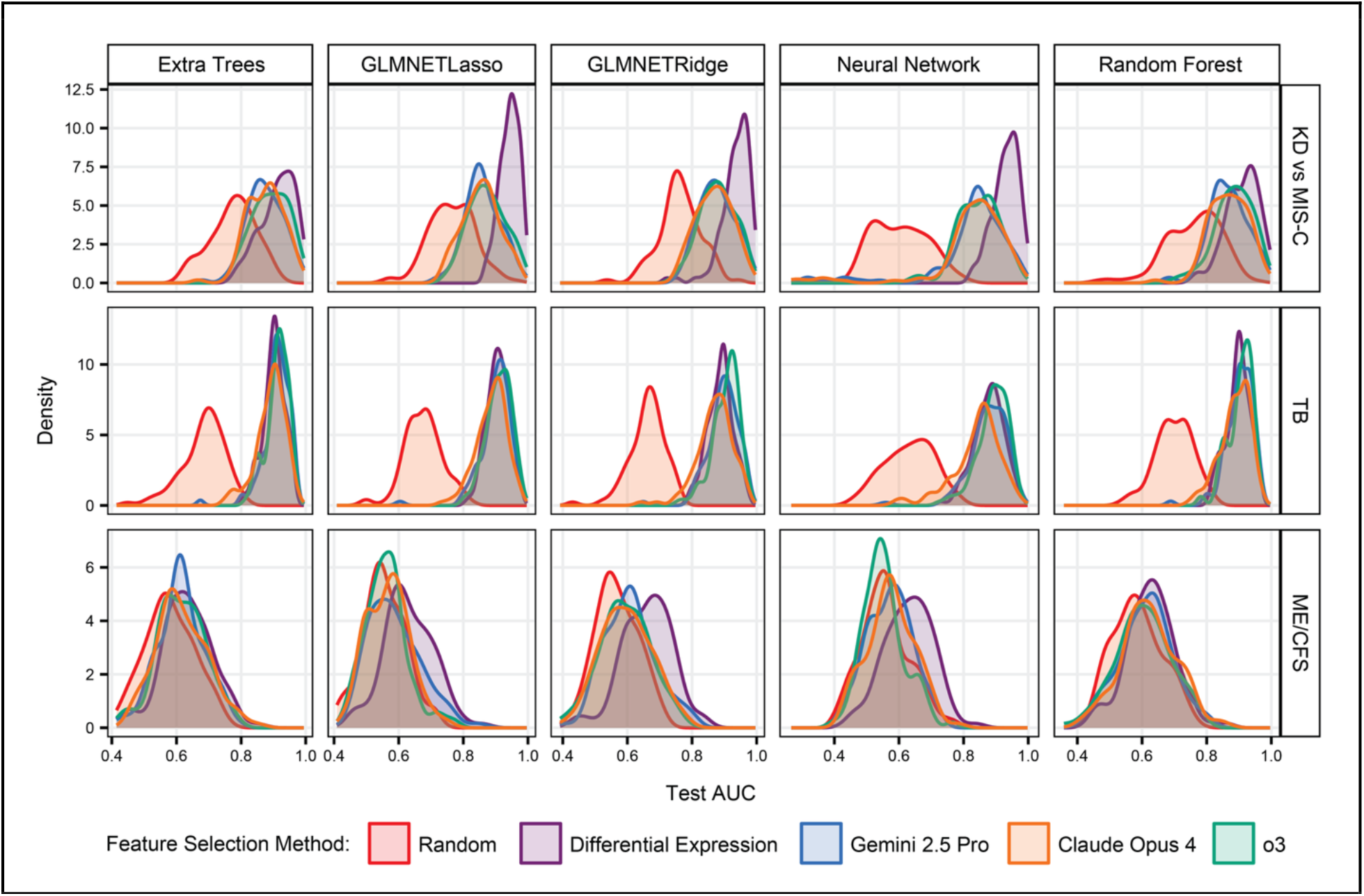
Distribution of ML ROC AUCs across 100 seeds using random gene selection, LLM gene selection, and DGE-based gene selection.

Together, these cohorts provide a spectrum, ranging from disorders with well-characterized biology to syndromes with no existing molecular tests, that serves as an ideal testing ground for evaluating LLMs. Depending on the task, each LLM received one or more of the following: (i) non-normalized plasma cfRNA read-count matrices (gene x sample), (ii) lists of gene names present in the read-count matrices (**File S1**), and (iii) curated text prompts of varying contextual depth (**Fig. 1B** and **Fig. 4A, *Methods***).

### LLMs vary in their ability to adhere to prompt requirements

Before assessing model performance, we evaluated each LLM’s ability to follow the prompt. For every cohort, we asked each LLM 100 times to draw exactly 200 genes from a reference list in order of predictive potential. We used either a “short” or “long” prompt. The short prompts (**Tables S1, S3, S5**) provided only minimal disease context, whereas the long prompts (**Tables S2, S4, S6**) offered guidance on redundancy reduction and specific approaches for utilizing evidence from the literature (***Methods***). Prompt adherence was quantified as the number of genes returned (**Fig. 1C and S1A**) and the percentage of selected genes present in the reference list (**Fig. 1D and S1B**). Claude 3.7 Sonnet and Claude Opus 4 showed comparable prompt adherence, whereas both OpenAI and Google demonstrated clear improvements in their newest model releases. Notably, Claude 3.7 Sonnet’s prompt adherence was on par with the newer OpenAI and Google models (**Files S2-S5**). GPT-4o and Gemini 2.0 Flash returned many non-reference features, including protein aliases (e.g., ASCT-1), endogenous retrovirus families (e.g., HERV-W), metabolites (e.g., NADPH), pathway labels (e.g., PI3K), non-human orthologues (e.g., SERPINB14), and even hallucinated features. Adherence and downstream predictive performance did not differ appreciably between short and long prompts, so all subsequent analyses used panels generated only with the long prompts.

### Performance of LLM-nominated gene panels

To test the predictive value of each gene panel, we used the first 100 genes from each iteration in a machine-learning pipeline (***Methods***). The pipeline trains five classifiers: ridge- and lasso-regularized generalized linear models (GLMNETRidge, GLMNETLasso), random forest, extra trees, and a simple feed-forward neural network. We benchmarked LLM-selected panels against panels based on DGE analysis and randomly selected genes. Because most of the initial panels selected by the LLMs included non-reference genes, many seeds included fewer than 100 features. Performance declined whenever a seed fell short of 100 usable genes or possessed genes expressed in only a few samples, due to a lack of informative candidate features for the models. For the KD vs. MIS-C cohort, across all classifier types, all LLM panels outperformed the random sets but underperformed the DGE panels with no significant differences between the LLMs. For the TB cohort, LLM panel performance matched the DGE panels, even exceeding them in mean receiver operating characteristic area under the curve (ROC AUC) values in many cases (o3 panels outperformed DGE panels across all five ML algorithms, Claude 3.7 Sonnet and Gemini 2.5 Pro across three), while outperforming the random sets across all five algorithms. Lastly, for the ME/CFS cohort, every LLM panel trailed the DGE panels and achieved only slight gains over random gene sets (**Fig. S2 and S3, Files S6 and S7**). Overall, performance relative to the DGE panels seemed to track the extent of knowledge in the public domain: TB, supported by abundant public data, reached high accuracy; ME/CFS, with a sparse knowledge base, lagged; KD vs. MIS-C fell in between.

### LLMs select disease-relevant genes

To investigate the biological relevance of the features selected by each model, we first tallied how often each gene appeared across the 100-gene panels and then analyzed the 20 most frequently chosen genes for each clinical cohort (**Fig. 3A and S4A**). In KD vs. MIS-C, the three LLMs converged strongly: ten genes were selected by all three models, eight of which (IL1B, IL6, STAT1, CXCL10, ICAM1, VCAM1, IL10, IRF7) were also significantly differentially expressed in the cfRNA data. TB and ME/CFS selections showed intermediate overlap across LLMs, with three genes selected by all three LLMs and DGE. For TB, o3 and Gemini 2.5 Pro selected GBP5, a well-validated marker with high statistical significance in the dataset^9^. Across all three top-20 lists (60 total), GPT-o3 and Gemini 2.5 Pro each captured 26 significantly expressed genes, whereas Claude Opus 4 captured 22. Claude Opus 4 selected the same gene across all three cohorts five times, compared with six times for GPT-o3 and Gemini 2.5 Pro. Only three features (IL1B, IL6, and TNF) appeared in every LLM’s top 20 list for every cohort, indicating condition specificity.

**Figure 3:**
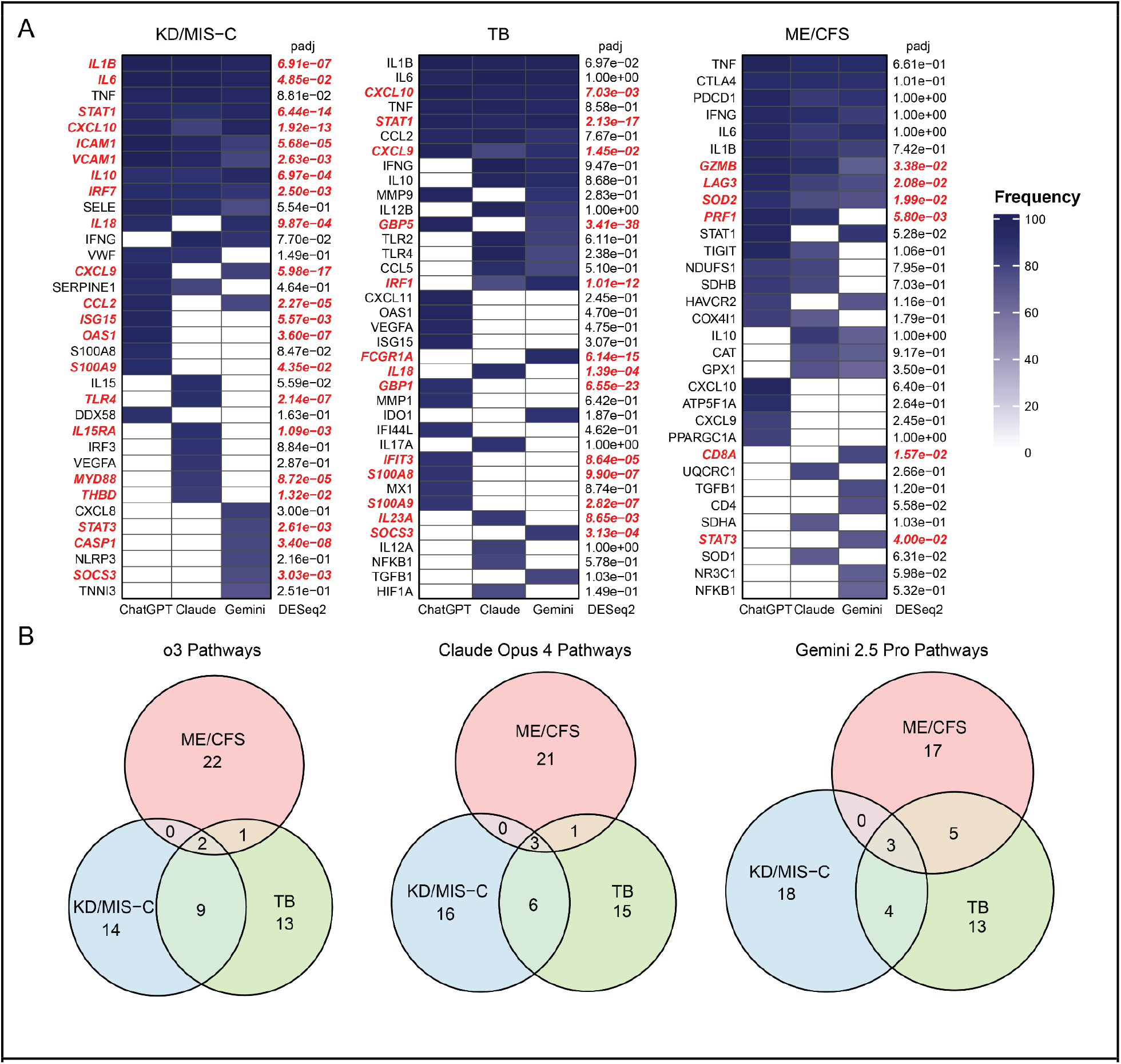
LLM gene selection and biological correlation. (A) Heatmap of the combined top 20 most frequently selected genes across LLMs and conditions with correspondence to expression in RNA-seq data (red = adjusted p value < 0.05), and (B) Venn diagram of top 25 enriched pathways based on LLM gene selection by condition.

Using selection frequency as a weight, we performed single-sample gene-set enrichment analysis (ssGSEA) and examined the 25 top-ranked pathways for each condition within each LLM (**Fig. 3B and S4B, File S8**). Each model enriched for immune and inflammatory pathways, such as IL-6, IL-17, and NF-κB signaling, yet the specific pathway mix differed by both LLM and disease. KD vs. MIS-C and TB showed the broadest consensus, whereas ME/CFS displayed the least overlap. This analysis shows that nominated genes were disease-related, and while all LLMs identified biologically plausible features across all cohorts, performance varied between models and with disease complexity, paralleling the trends seen in classifier accuracy.

### LLMs construct classifiers with high accuracy

After benchmarking gene feature selection by LLMs, we tested whether the LLMs could perform the entire ML workflow autonomously without intervention. Starting with a cfRNA count matrix and corresponding diagnosis labels for the training set, we instructed the LLM to select features and construct and train a binary classifier. We then provided a test set without annotation and instructed the model to classify each sample. The workflow was executed 50 times with disease-naïve prompts, omitting any information related to condition, cfRNA, sample type, or diagnosis (**Table S7**), and 50 times with disease-informed prompts, which provided details on the condition and sample type and encouraged the LLM to utilize internal or external domain knowledge (**Tables S8–S10**). In parallel, we performed 50 bootstrap iterations of the standard ML pipeline (**Fig. 4A**).

**Figure 4:**
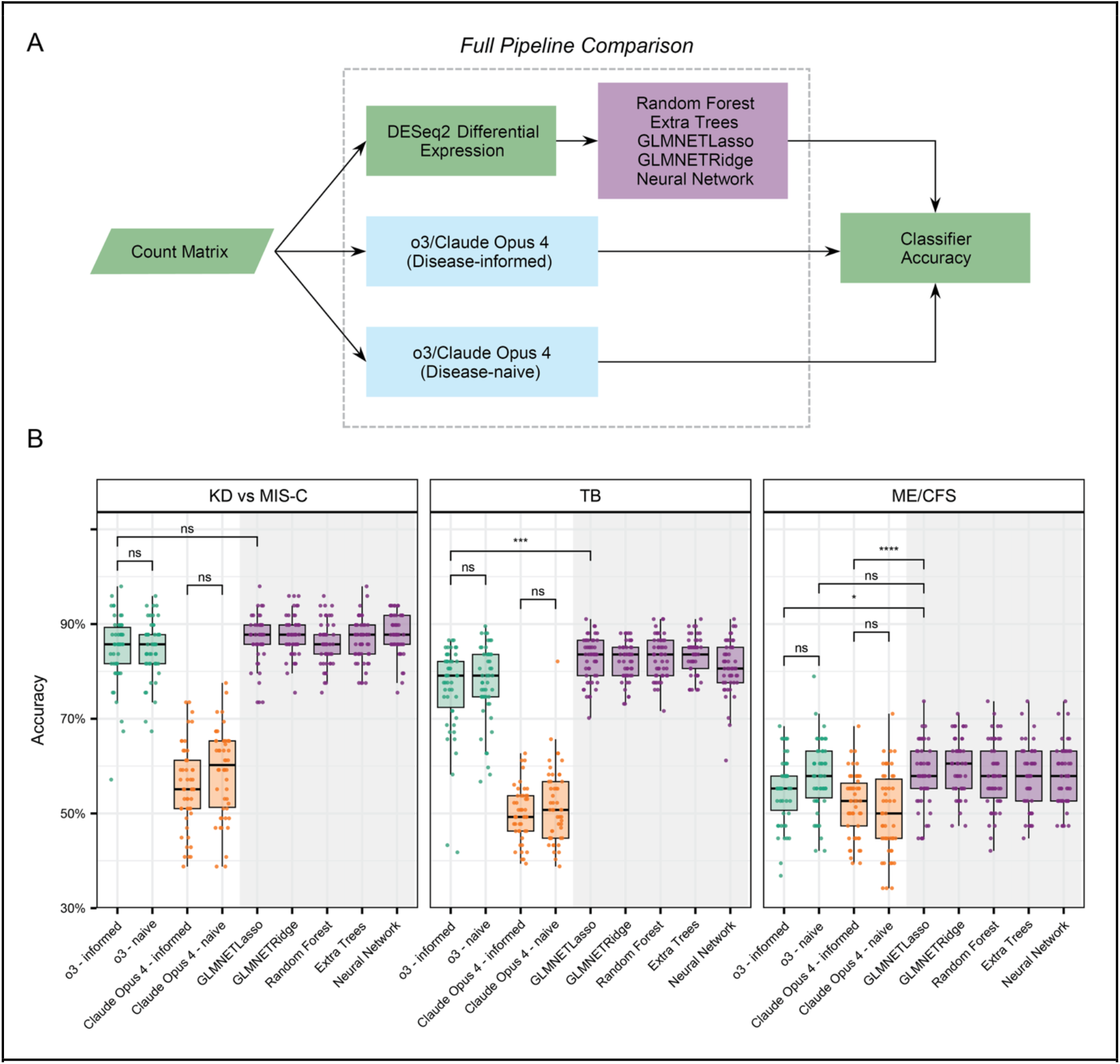
LLM-constructed classifier performance. (A) Workflow overview for comparison between LLM and ML classifiers and (B) accuracy distributions for ML, disease-informed LLM, and disease-naïve LLM classifiers.

Among the most advanced models we evaluated (o3, Claude Opus 4, Gemini 2.5 Pro), only o3 and Claude Opus 4 were capable of consistently performing this task. Gemini 2.5 Pro typically remained in an indefinite “thinking” state or terminated with runtime errors. In the KD vs. MIS-C cohort, o3 produced classifiers with mean accuracy approaching that of the GLMNETLasso benchmark (mean=86.7%), with virtually identical results with disease-informed (84.7%) and disease-naïve (84.8%) prompts (**Fig. 4B)**. Claude Opus 4 performed significantly worse than both o3 and the benchmark with either prompt type. In the TB cohort, the performance gap widened. The standard workflow reached 82.6% mean accuracy, whereas o3 achieved 76.3% with disease-informed and 77.6% with disease-naïve prompts. Claude Opus 4 again trailed with disease-informed and disease-naïve prompts accuracies of 49.9% and 51.6%, respectively. For the ME/CFS cohort, all methods struggled to classify with high accuracy. Disease-naïve o3 runs achieved 57.7% accuracy, the only LLM condition not statistically inferior to the benchmark (57.9%), while disease-informed o3 (55.0%), disease-informed Claude Opus 4 (52.1%), and disease-naïve Claude Opus 4 (50.0%) performed worse (all adjusted p < 0.05). Across all three clinical cohorts, providing clinical context (disease-informed vs. disease-naïve prompts) did not improve accuracy (KD vs. MIS-C: o3 p=0.890, Claude Opus 4 p=0.147; TB: o3 p=0.445, Claude Opus 4 p=0.249; ME/CFS: o3 p=0.113, Claude Opus 4 p=0.230) (**Files S9 and S10**). Of note, when given disease-informed prompts, both o3 and Claude Opus 4 created and used pathway-level variables as features for ML (e.g., “pathway_INTERFERON_GAMMA” and “Pathway_IL1_TNF_score”), but the addition of these features did not improve the performance.

### User experience

For LLM-driven feature selection, when given the prompt and reference gene list, each LLM returned ∼200 features within ≈ 2 min. Failures clustered into two categories: (i) refusals citing missing metadata and (ii) sessions that “lost’’ the uploaded file. Failures were more frequent with Google models than with OpenAI or Anthropic models. For the end-to-end pipeline task, only o3 and Claude Opus 4 could consistently complete both training and inference; Gemini 2.5 Pro timed out or refused execution and was dropped from further analysis. o3 trained in 4–6 min and inferred in 3–5 min, with occasional file-access errors. Claude Opus 4 trained more slowly (8–11 min) and failed more often. Importantly, none of the models produced code that ran unmodified outside the chat environment. Extracted code snippets required substantial manual integration, limiting reproducibility.

## DISCUSSION

We assessed the utility of LLMs for liquid-biopsy biomarker discovery. Despite prompt-adherence issues, all LLMs recovered biologically meaningful signals from plasma cfRNA-seq data. Convergence was strongest in the KD vs. MIS-C and TB cohorts and weakest for ME/CFS, a condition with limited mechanistic consensus and sparse literature. Gene-abundance weighting and pathway enrichment showed that each model converged on canonical immune pathways: interleukin signaling in KD vs. MIS-C, NK-cell and interferon signaling in TB, and mitochondrial stress, interferon, and interleukin modules in ME/CFS.

We compared ML classifiers trained using gene panels selected at random, by the LLMs, and with DGE analysis. Across clinical cohorts, the LLM-nominated gene panels outperformed the randomly selected gene panels, matched DGE panels in TB, but under-performed DGE in KD vs.

MIS-C and ME/CFS. A considerable performance increase was observed between older and newer LLMs from OpenAI and Google, highlighting the rapid improvements being made between LLM generations. OpenAI advanced its flagship model in ∼11 months (May 2024 to April 2025) and Google in ∼3 months (December 2024 to March 2025). Notably, the older Claude 3.7 Sonnet still matched or exceeded the newer OpenAI and Google models in prompt adherence and performance of this task.

A different picture emerged when the LLMs were asked to build complete workflows from raw sequencing counts. o3 generated end-to-end classifiers for KD vs. MIS-C that matched the performance of the standard workflow (naïve p=0.137, informed p=0.137), but underperformed the standard workflow in TB (naïve p=2.470e-05, informed p=5.299e-05) and offered no improvement for ME/CFS (naïve p=0.966, informed p=0.066). Providing additional disease context did not improve accuracy, implying that performance gains stemmed from code-generation rather than domain knowledge.

Collectively, we find that current LLMs can extract biologically meaningful gene candidates and automate large portions of a bioinformatics pipeline, yet traditional statistical or hybrid workflows still deliver the highest predictive performance in most cases. As LLMs improve, benchmarks like this one will be critical for transparent, reproducible, and clinically safe deployment of language-model diagnostics, shifting the bottleneck from panel discovery to rigorous validation and clinical implementation of AI-generated disease signatures.

## METHODS

### Sample acquisition and data pre-processing

Sample acquisition, clinical diagnosis information, sample processing and sequencing, and sample quality filtering for KD and MIS-C patients^7,27^, TB-positive and TB-negative patients^9,28–30^, and ME/CFS patients and sedentary controls^11,31^ has been previously described.

### Sample partitioning

Samples were partitioned into training and test sets at a ratio of 70:30. TB cohort samples were partitioned for ML applications evenly based on collection location and sex. Sample partitioning of the ME/CFS cohort took into consideration collection location, sex, and sequencing batch information.

### In-house machine learning pipeline development

After partitioning and seed-splitting, differential abundance analysis was conducted on the training data. Genes were filtered based on specific criteria for each clinical cohort (KD vs. MIS-C: adjusted *P*-value < 0.05, base mean abundance > 50, and absolute log2 fold change > 1; TB: adjusted *P*-value < 0.05 and base mean abundance > 50; ME/CFS: *P*-value < 0.001 and base mean abundance > 50). Filtering criteria and threshold testing have been previously explained^7^. Raw counts for the training and test sets were individually normalized with variance stabilizing transformation using the DESeq2 package. The test set was transformed using the dispersion function from the training set. Five ML classification algorithms were employed using the R package Caret^41^, including generalized linear models with ridge-or lasso-regularization (GLMNETRidge, GLMNETLasso), random forest, extra-trees ensemble, and a feed-forward neural network. Training was performed using fivefold cross-validation and grid search hyperparameter tuning. After training each model, we fixed a decision threshold by maximizing Youden’s J statistic on the training set. We then applied that threshold to the held-out test samples to generate class labels and reported overall classification accuracy as the performance metric.

### Open-domain gene panel selection

#### LLM input prompt generation

To demonstrate response variability due to input prompt quality, two input prompts were designed for each clinical cohort. The “short” prompts (**Tables S1, S3, and S5**) only stated the two conditions within each cohort, providing no additional insight or guidance on how to approach gene selection. The “long” prompts (**Tables S2, S4, and S6**), however, included information on biological pathway relevance and suggested considerations for handling literature and database evidence, minimizing pathway redundancy, and optimizing diagnostic potential of the selected gene panel. For the KD vs. MIS-C and TB cohorts, the LLMs were instructed to avoid using information from peer-reviewed articles previously published by our lab that used an overlapping dataset and were available online during the time of data generation.

#### LLM querying

Six proprietary LLMs (OpenAI’s o3 and GPT-4o, Google’s Gemini 2.5 Po and Gemini 2.0 Flash, and Anthropic’s Claude Opus 4 and Claude 3.7 Sonnet) were prompted to select genes for a diagnostic panel from a provided reference list. This list, attached in the input prompt as a CSV file, contained all genes from the RNA-seq expression matrices that passed quality control filtering. For each clinical cohort, each input prompt and paired gene list was submitted to each LLM through their respective web interfaces 100 times, generating 100 individual gene panels per prompt type, per LLM, per cohort. This allowed for the assessment of output reproducibility and consistency across models and prompts. The resulting distinct feature sets were then assigned to individual seeds and evaluated in a downstream ML pipeline. The “share-memory” (conversation-history) feature was turned off; each prompt was submitted in a fresh session, eliminating cross-conversation context.

#### LLM-selected gene panel evaluation: input prompt adherence

Each set of LLM-selected gene panels were evaluated for their adherence to the requirements of the input prompt. Specifically, the resulting panel should contain exactly 200 genes, all of which should be found in the provided list. Deviation from 200 genes and the percentage of genes not present on the provided reference list were calculated for each panel. Statistical differences in input prompt adherence between gene panels were evaluated with a two-tailed paired t-test.

#### LLM-selected gene panel evaluation: gene and enriched pathway analysis

Count matrices of gene selection were generated for each condition and LLM. With the number of LLM selections being used as an analogue for gene count, these matrices were analyzed using QIAGEN Ingenuity Pathway Analysis (IPA)^42^ to identify the top enriched pathways, providing a measure of the biological plausibility of LLM-driven gene selection.

#### LLM-selected gene panel evaluation: predictive performance benchmarking with machine learning

LLM-selected gene panels were benchmarked against both randomly generated panels and panels derived from DGE analysis using a ML evaluation framework. DGE analysis was performed using the DESeq2 R package^43,44^. For each feature set, 100 genes were selected. For LLM generated features we chose to select the first 100 ranked genes from the 200 available as opposed to requesting that the LLMs return 100 genes, as prompt adherence limitations would make many of the seeds unusable for predictive modeling. In the KD vs. MIS-C cohort, genes with a base mean transcript abundance >50 and/or absolute log_2_ fold change >1 were excluded, and the top 100 genes with the lowest adjusted *P*-values were selected. In the TB and ME/CFS cohorts, genes with base mean transcript abundance > 50 were filtered out, followed by selection of the top 100 genes based on adjusted *P*-value (TB) or unadjusted *P*-value (ME/CFS). For each clinical cohort, 100 random seeds were generated, and samples were partitioned into training and testing sets using a 70:30 split, with cohort-specific constraints applied (***Methods: Sample partitioning***). Feature sets were fed into the in-house ML pipeline and final model performances were evaluated on the held-out test set using ROC-AUC values.

### Predictive pipeline development

#### LLM input prompt generation

To evaluate the LLM’s capacity to integrate domain knowledge into pipeline development, we generated two prompt types for each clinical cohort: “disease-naïve” (**Table S7**) and “disease-informed” prompts (**Tables S8-S10**). The disease-naïve prompt instructed the LLM to construct a binary classifier without referencing cfRNA, disease context, or related biological concepts. In contrast, the disease-informed prompt explicitly described the input data as plasma-derived cfRNA count matrices and specified the disease phenotypes and control groups for each classification task. The disease-informed prompt encouraged utilization of both parametric knowledge (encoded during training) and non-parametric knowledge (external retrieval), while explicitly excluding relevant peer-reviewed publications from our research group to prevent data leakage. Both prompt configurations specified identical output requirements: (i) notification upon training completion with a request for held-out test data, (ii) generation of predicted class labels for all test samples, and (iii) output of all non-zero weighted features ranked by absolute weight magnitude. Additionally, the LLM was instructed to indicate which features were informed by prior domain knowledge, enabling identification of parametric and non-parametric knowledge contributions to feature selection.

#### LLM querying

Three proprietary LLMs were evaluated for their ability to develop binary classifiers: OpenAI’s o3, Google’s Gemini 2.5 Pro, and Anthropic’s Claude Opus 4. Each model was tasked with differentiating disease states within each clinical cohort using either disease-naïve or disease-informed prompts. Only o3 and Claude Opus 4 were able to consistently and successfully complete the full pipeline, including model development and prediction generation on held-out test data. The experimental workflow proceeded as follows: the LLM received an input prompt (disease-naïve or disease-informed) through the web interface along with training data in CSV format. Upon completion of binary classifier construction, the LLM requested provision of held-out test data (CSV format). Training and testing dataset preparation followed protocols detailed in ***Methods: Sample acquisition and data pre-processing*** and ***Methods: Sample partitioning*** sections. Subsequently, the LLM generated two CSV output files as specified in the input prompt: predicted class labels for test samples and ranked non-zero weighted features. These output files were directly downloaded from the interface for downstream analysis. This protocol was executed 50 times per clinical cohort for both prompt conditions (disease-naïve and disease-informed), yielding 100 total runs per cohort per LLM. Each iteration employed unique, randomly generated seeds for training-testing partitioning, with identical seeds maintained across all comparative ML analyses to ensure consistency. The “share-memory” (conversation-history) feature was turned off; each prompt was submitted in a fresh session, eliminating cross-conversation context.

#### LLM-generated predictive pipeline evaluation: predictive accuracy

Accuracy values for each seed were generated by comparing the LLM-predicted class to the true class. Accuracy values generated by the LLM for each prompt-type were then plotted against accuracy values from the standard pipeline. Differences in classifier performance were assessed with two-tailed paired t-tests when both models were evaluated on the same samples and two-tailed Welch’s t-tests when the sample sets differed.

## DATA AVAILABILITY

The data supporting the findings of this study, including the summary statistics underlying the figures and tables, are available within the paper and its ***Supplementary Information*** files.

## CODE AVAILABILITY

All custom code used for data processing, analysis, and figure generation is available on GitHub (https://github.com/adb258/cfrna_ai_manuscript).

## ETHICS STATEMENT

The Cornell University IRB for Human Participants (2012010003), New York, NY, approved the protocols for this study. All samples and patient information were deidentified for analysis and shared with collaborating institutions.

## AUTHOR CONTRIBUTIONS

H.A.G., A.B., C.J.L., and I.D.V. designed research; H.A.G., A.B., C.J.L., D.E.L., A.E.G., and I.D.V. analyzed data; H.A.G., A.B., and I.D.V. wrote the paper.

